# Milk-Way Algorithm applied in Imbalanced Dataset

**DOI:** 10.1101/2021.02.22.432316

**Authors:** Carmelina Figueiredo Vieira Leite, Marcos Augusto dos Santos

**Author notes:** Equal contribution.

## Abstract

We wish to evaluate the algorithm Milk-Way, using a known dataset deposited in a public repository. The new algorithm, which converges various techniques from different areas of knowledge, can classify ligands and select potential new drugs. It was used a dataset of ligands, organized by 15 Bioassays and described by different fingerprints. Full details of the dataset architecture were already published in a public repository. Through the stratified feature selection, using the Milk-Way algorithm, the True Positive and False Positive Rates reached a higher performance compared to the published paper. Using all the features available for each Bioassay, we reached the lowest metrics in all of them. We demonstrated that adding more features have not made a significant impact on the performance. In all the Bioassays, the True Positives and False Positives reached 100% and 0%, respectively, only using 50% and 75% of the features available. The Milk-Way algorithm suggests a holistic approach, which will contribute to the machine-learning area, namely to classified ligands in the virtual screening.

## Background

Small-molecule drugs, which are the constituents of most therapies, can be defined as non-polymeric low-molecular-mass (<1000 Da) organic compounds (1). High-throughput screening (HTS) is a common first step to discover new chemical structures (2–4). Within a few weeks, HTS enables testing 1−5 million compounds (5), still being an inadequate sample due to the high number of possible drug molecules existing in the chemical space (2–4). The goal of HTS is to find new compounds for chemical optimization. Therefore, false-positives may be tolerated as long as leads are truly found (3, 4, 6). These molecules reacted positively in an assay that operates via irrelevant mechanisms (3, 6). One of the reasons for a high number of false-positives is promiscuity (3).

PubChem (http://pubchem.ncbi.nlm.nih.gov) is an open public repository containing chemical structures and biological properties of molecules including small molecules, siRNAs, miRNAs, carbohydrates, lipids, peptides, chemically modified macromolecules and many others, hosted by the National Center for Biotechnology Information since 2004 (7–9).

Three interconnected databases constitute the PubChem: Substance, BioAssay and Compound. The first database contains contributed sample descriptions (primarily small molecules) provided by depositors. The Compound database contains the unique chemical structures derived from the Substance database records (8, 9). Each BioAssay record, provided by depositors, can contain as many comments and as much descriptive text as needed to provide the assay’s overall background. This can include the biological system tested in the assay, the relationship between a disease and the selected therapeutic target (8). The label to identify Substance, Compound and BioAssay databases are SID (SubstanceID), CID (CompoundID) and AID (AssayID), respectively (9).

The BioAssay database has been growing substantially during the past years (8). In September of 2013, the BioAssay database has received >700000 depositions of bioassays. This represents thousands of potential modulators for >8000 protein targets (7). In September 2015, it had more than 157 million depositor-provided chemical substance descriptions, covering about 10 thousand unique protein target sequences (9).

The stage of the biological experiments in PubChem varies. These methods include: ‘screening’ assay, usually a primary high-throughput assay where the activity outcome is based on a percentage inhibition from a single dose; ‘confirmatory’ assay, typically a low-throughput assay where the activity outcome is based on a dose–response relationship with multiple tested concentrations (8). Because of this, it is possible to find substance records that are active in a primary screening assay but inactive in a corresponding confirmatory assay.

The datasets deposited in PubChem are highly imbalanced, with the ratio of active to inactive compounds on an average of 1:1,000. He points out that the false-positive rates in primary screens are high, although many actives are not confirmed even in secondary and/or confirmatory assays (10). It becomes necessary to apply techniques that allow the suitable extraction of data.

There are informatics techniques, which have been developed to shorten the research cycle and reduce the expense and risk of failure for drug discovery (1, 11, 12). A decision tree (5, 13, 14), neural network (15, 16) and support vector machine (17) are examples of classification models that are constructed on an adequate and relatively balanced set of training data to draw an estimated decision amongst the different classes (17). Despite the techniques to minimize these difficulties (17, 18), there are still challenges to overcome (17).

In this paper, we built models for virtual screening of bioassays, using an available dataset (19, 20), which had already been used in a reclassification study through conventional methods. The method used was an adaptation of the Milk-Way algorithm (MW) (21, 22), which exceeded the performance of the results published.

## Method

### Data Collection

We used the data of Schierz (19), also available in UCI repository (20). A selection of Weka cost-sensitive classifiers was applied: Naive Bayes (NB), Support Vector Machine (SVM), J48.5 and Random Forest (RF) were implemented to a variety of bioassay datasets.

### Algorithm and Validation

We used the Milk-Way algorithm (21, 22) and applied the *k*-cross-validation (*k*=5) to evaluate the method’s performance.

## Results

We used a dataset from a paper already published with Bioassays, which employed a conventional classifier of data mining methods: NB, RF, SVM and J48 (19). In the present work, we use the same dataset available by Schierz (19) and compare the performance of both using the same measures – True Positive Rate (TP%) and False Positive Rate (FP%).

The True positive rate measures the proportion of correct predictions of active drugs compared to all positive samples, which are classified as actives and are calculated as follows:

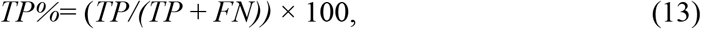

where TP is the number of correct predictions (true positive), while FN is the number of incorrect predictions (false negative) (23, 24). In our study, true positive means that it was in fact an active drug.

The False positive rate measures the proportion of the number of incorrect predictions compared to the whole number of truly inactive drugs. It is calculated as follows:

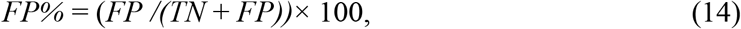

where TN is the number of correctly inactive samples (true negative) while FP is the number of incorrectly active samples (false positive). In our study, true positive means that it was not an active drug. These two parameters are associated with prediction error, which is 100% sensitive and specific, consequently being a good predictor (23, 24).

We evaluated all models with different metrics: sensitivity, specificity, precision, f1, fa, accuracy and AUC (See **Supplementary Material_Metrics**). All P(*x*) values obtained are also available (See **Supplementary Material_Values**). For example, we provide the dataset subjected to stratified feature selection of 5% (see **Supplementary Material_Dataset_5FS**).

All the data was processed in MATLAB R2019b, using a laptop with 8 GB RAM, 551 GB hard drive, and a processor Intel Core i7 2.70GHz.

**Table 1** shows the results of the MW algorithm, using the stratified feature selection (21), with the different % of the features used. When we write 100%, it corresponds to the total features that the dataset made available to us. For example, in AID362, 100% corresponds to 144 features; 75% to 108, 50% to 72, and 5% to 7 features. Unfortunately, with the MW, it was not possible to use the AID 1608 because the label in the dataset was active and inconclusive. In the Primary Bioassays, and with the Mixed Primary Screen/Confirmatory Bioassays, there is no variation with the aggregation of new features, reaching the point of worsening its performance using all the datasets’ features.

**Table 1.**
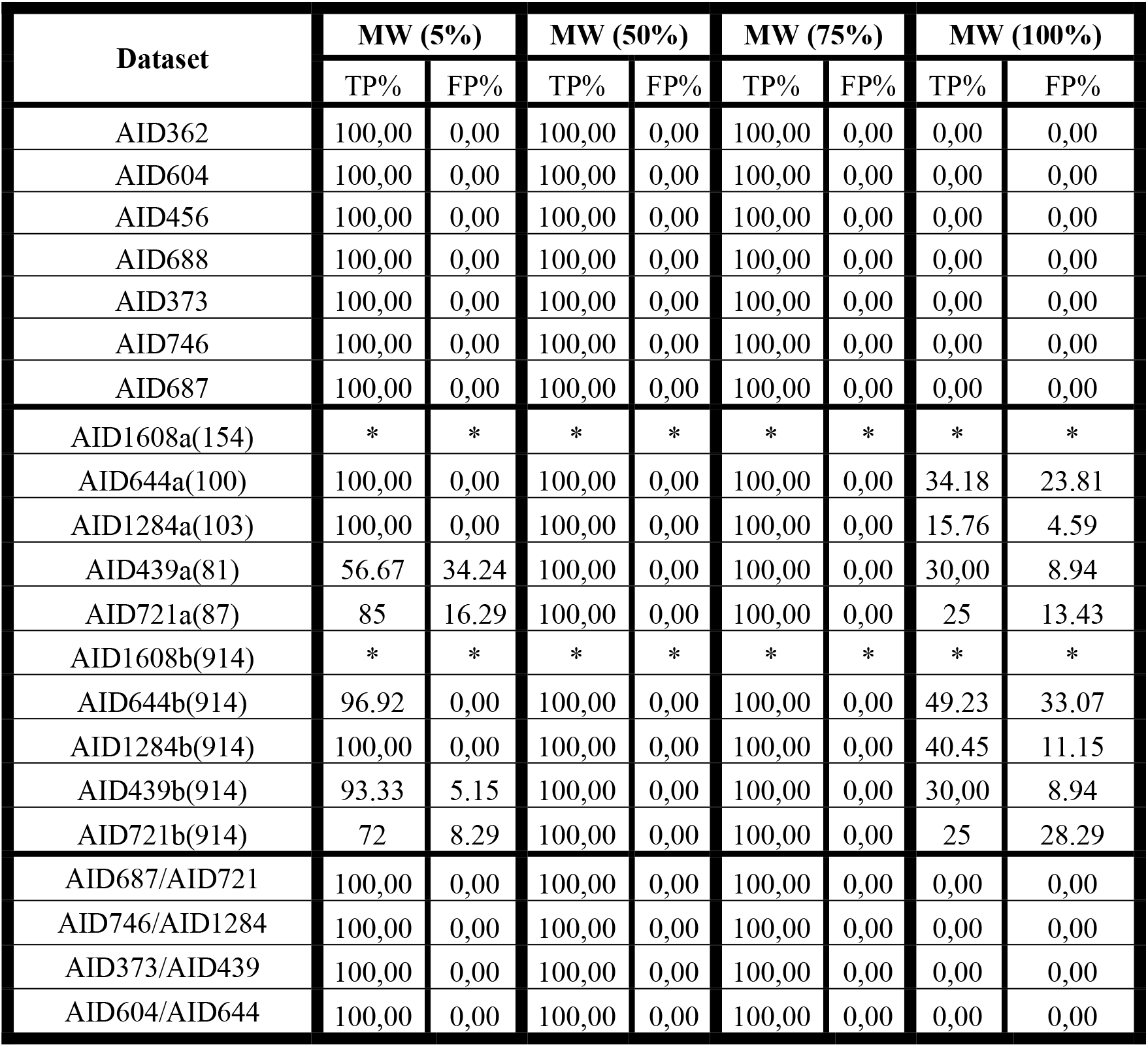
Results of all Bioassays using MW with different % of features used (*not possible to calculate)

| Dataset | MW (5%) |  | MW (50%) |  | MW (75%) |  | MW (100%) |  |
| --- | --- | --- | --- | --- | --- | --- | --- | --- |
|  | TP% | FP% | TP% | FP% | TP% | FP% | TP% | FP% |
| AID362 | 100,00 | 0,00 | 100,00 | 0,00 | 100,00 | 0,00 | 0,00 | 0,00 |
| AID604 | 100,00 | 0,00 | 100,00 | 0,00 | 100,00 | 0,00 | 0,00 | 0,00 |
| AID456 | 100,00 | 0,00 | 100,00 | 0,00 | 100,00 | 0,00 | 0,00 | 0,00 |
| AID688 | 100,00 | 0,00 | 100,00 | 0,00 | 100,00 | 0,00 | 0,00 | 0,00 |
| AID373 | 100,00 | 0,00 | 100,00 | 0,00 | 100,00 | 0,00 | 0,00 | 0,00 |
| AID746 | 100,00 | 0,00 | 100,00 | 0,00 | 100,00 | 0,00 | 0,00 | 0,00 |
| AID687 | 100,00 | 0,00 | 100,00 | 0,00 | 100,00 | 0,00 | 0,00 | 0,00 |
| AID1608a(154) | * | * | * | * | * | * | * | * |
| AID644a(100) | 100,00 | 0,00 | 100,00 | 0,00 | 100,00 | 0,00 | 34.18 | 23.81 |
| AID1284a(103) | 100,00 | 0,00 | 100,00 | 0,00 | 100,00 | 0,00 | 15.76 | 4.59 |
| AID439a(81) | 56.67 | 34.24 | 100,00 | 0,00 | 100,00 | 0,00 | 30,00 | 8.94 |
| AID721a(87) | 85 | 16.29 | 100,00 | 0,00 | 100,00 | 0,00 | 25 | 13.43 |
| AID1608b(914) | * | * | * | * | * | * | * | * |
| AID644b(914) | 96.92 | 0,00 | 100,00 | 0,00 | 100,00 | 0,00 | 49.23 | 33.07 |
| AID1284b(914) | 100,00 | 0,00 | 100,00 | 0,00 | 100,00 | 0,00 | 40.45 | 11.15 |
| AID439b(914) | 93.33 | 5.15 | 100,00 | 0,00 | 100,00 | 0,00 | 30,00 | 8.94 |
| AID721b(914) | 72 | 8.29 | 100,00 | 0,00 | 100,00 | 0,00 | 25 | 28.29 |
| AID687/AID721 | 100,00 | 0,00 | 100,00 | 0,00 | 100,00 | 0,00 | 0,00 | 0,00 |
| AID746/AID1284 | 100,00 | 0,00 | 100,00 | 0,00 | 100,00 | 0,00 | 0,00 | 0,00 |
| AID373/AID439 | 100,00 | 0,00 | 100,00 | 0,00 | 100,00 | 0,00 | 0,00 | 0,00 |
| AID604/AID644 | 100,00 | 0,00 | 100,00 | 0,00 | 100,00 | 0,00 | 0,00 | 0,00 |

The primary Bioassay (**Table 2**), and combining these with the confirmatory ones (**Table 4**), demonstrates that the performance is superior using only 5% of the features than when using the MW at 100% and all the algorithms applied.

**Table 2.**
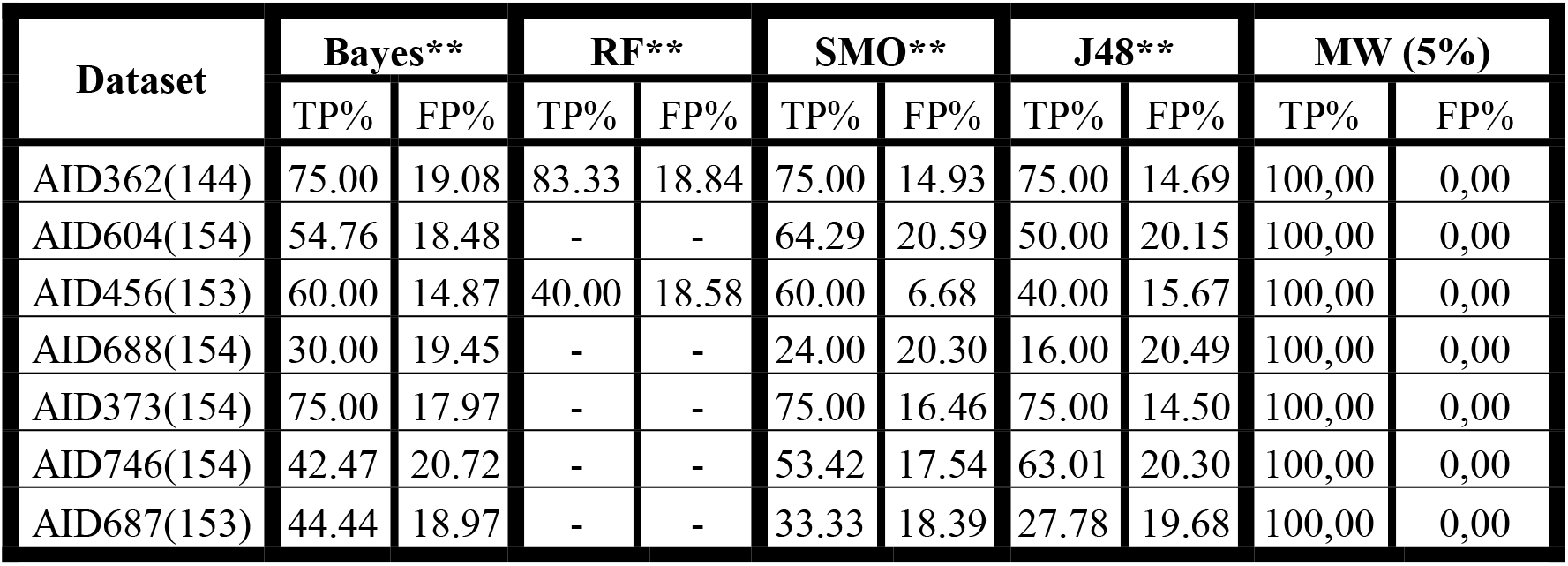
Results of all methods, with primary Bioassays. **available in Schierz (2009).

With confirmatory Biossays (**Table 3**), which are more specific datasets, the performance variation is more noticeable. When we use all the features, 100%, AID 644 (b) and AID1284 (b) increase both TP% and FP%. AID439 (b) and AID721 (b) only increase FP. However, if we use only 5%, all TP% are above 72% and FP% below 8.29. The selection of 50% and 75% of the features is when they all reach 100% in TP and FP. The values of P(*x*) (See **Supplementary Material_Values)** are very similar, which gives us the indication that alphas will be very similar too. The cases of AID 721(b) and 439(b) are flagrant. Alpha values of AID721(b) using 75% (686 features) only added alphas equal to zero, relating with AID 721(b) using 50% (457 features) (See **Table 2** and **Supplementary Material_Alphas).** The sum has no meaning, it just shows that zeros are added.

**Table 3.** Results of all methods, with confirmatory Bioassays. * Inconclusive Bioassays can not be used in modified logistic regression.**available in Schierz (2009).

| Dataset | Bayes** |  | SMO** |  | RF** |  | J48** |  | MW (5%) |  |
| --- | --- | --- | --- | --- | --- | --- | --- | --- | --- | --- |
|  | TP% | FP% | TP% | FP% | TP% | FP% | TP% | FP% | TP% | FP% |
| AID1608a(154) | 23.08 | 19.17 | 30.77 | 8.81 | 30.77 | 8.29 | 15.78 | 20.21 | * | * |
| AID644a(100) | 38.46 | 39.29 | 23.08 | 17.86 | 23.08 | 7.14 | 38.46 | 28.57 | 100.00 | 0.00 |
| AID1284a(103) | 27.27 | 26.23 | 36.36 | 13.11 | 45.45 | 18.03 | 54.55 | 13.11 | 100.00 | 0.00 |
| AID439a(81) | 100.00 | 27.27 | 50.00 | 9.09 | 50.00 | 18.18 | 50.00 | 18.18 | 56.67 | 34.24 |
| AID721a(87) | 0.00 | 28.57 | 0.00 | 14.29 | 0.00 | 21.43 | 0.00 | 14.29 | 85.00 | 16.29 |
| AID1608b(914) | 30.77 | 12.95 | 38.46 | 13.47 | 30.77 | 12.95 | 38.46 | 18.13 | * | * |
| AID644b(914) | 30.77 | 28.57 | 53.85 | 14.29 | 38.46 | 17.86 | 30.77 | 14.29 | 96.92 | 0.00 |
| AID1284b(914) | 36.36 | 24.59 | 36.36 | 13.11 | 54.55 | 18.03 | 45.45 | 16.39 | 100.00 | 0.00 |
| AID439b(914) | 50.00 | 18.18 | 50.00 | 9.09 | 50.00 | 18.18 | 0.00 | 27.27 | 93.33 | 5.15 |
| AID721b(914) | 0.00 | 14.29 | 0.00 | 21.43 | 0.00 | 7.14 | 0.00 | 0.00 | 72.00 | 8.29 |

**Table 4.** Results of all methods, with primary/confirmatory Bioassays. * out of memory.**available in Schierz (2009).

| Dataset | NB** |  | RF** |  | SMO** |  | J48** |  | MW(5%) |  |
| --- | --- | --- | --- | --- | --- | --- | --- | --- | --- | --- |
|  | TP% | FP% | TP% | FP% | TP% | FP% | TP% | FP% | TP% | FP% |
| AID687/AID721 | 50.00 | 14.56 | - | - | 50.00 | 22.53 | 50.00 | 9.49 | 100,00 | 0,00 |
| AID746/AID1284 | 72.73 | 17.70 | - | - | 81.82 | 14.98 | 100.00 | 19.34 | 100,00 | 0,00 |
| AID373/AID439 | 50.00 | 2.33 | - | - | 50.00 | 8.86 | 100.00 | 17.65 | 100,00 | 0,00 |
| AID604/AID644 | 46.15 | 12.91 | - | - | 76.92 | 11.91 | 53.85 | 12.17 | 100,00 | 0,00 |

**Table 5.** Sum of alpha values of each fold (k-cross-validation, k=5).

| Dataset | AID439 |  | AID721 |  |
| --- | --- | --- | --- | --- |
| Features | 81 | 914 | 87 | 914 |
| MW (5%)<br>(sum of alpha values<br>of each fold) | -6.140226918 | -46.59887642 | -20.32345719 | 9.233761835 |
|  | -4.605170191 | -33.5790307 | -23.99967905 | 17.7768792 |
|  | -9.518820673 | -37.68413048 | -34.11082261 | 6.524484158 |
|  | -12.45146811 | -52.87246571 | -23.0240244 | -12.67241821 |
|  | -2.763102118 | -46.49814292 | -22.05936116 | -1.357405339 |
| MW (50%)<br>(sum of alpha values<br>of each fold) | -0.231240509 | 4.735433129 | -4.89563565 | 3.136741595 |
|  | 0.406994687 | 6.433720229 | -0.806924038 | 3.946762375 |
|  | 0.857566961 | 4.380051342 | 0.6345217 | 1.839487346 |
|  | -1.000767464 | 5.056561762 | -0.098936266 | 7.129050638 |
|  | 2.501519829 | 7.939542566 | 1.188224216 | 4.788912727 |
| MW (75%)<br>(sum of alpha values<br>of each fold) | -1.869919141 | 4.735433129 | 0.033305591 | 3.136741595 |
|  | -3.497116231 | 6.433720229 | 2.269059114 | 3.946762375 |
|  | -3.167833375 | 4.380051342 | 2.309428311 | 1.839487346 |
|  | -4.223065441 | 5.056561762 | 2.547239151 | 7.129050638 |
|  | -0.946482813 | 7.939542566 | 4.385760849 | 4.788912727 |
| MW (100%)<br>(sum of alpha values<br>of each fold) | 53.03307114 | 11.03625221 | 19.52887105 | 41.36371797 |
|  | 60.07973628 | 24.49394849 | 4.369732777 | -14.79235448 |
|  | 69.6363323 | -2.822035056 | -28.54330355 | 17.01549603 |
|  | 69.53548713 | 14.19361035 | 19.9733208 | 3.680885492 |
|  | 69.57739905 | 5.400712047 | 29.64185058 | 60.28832346 |

Comparing the paper with the available dataset (19), both TP% and FP% exceed. It is worth acknowledging that in the original paper, the author had taken some features, indicating different numbers for each Bioassay. This forecasts that using too many features does not help the performance.

## Discussion and Conclusion

Submitting an entire compound library to a molecular profiling panel can easily become cost-prohibitive. Using PubChem HTS fingerprints as molecular descriptors shown to be efficient in retrieving hits that are structurally diverse and distinct from active compounds retrieved by chemical similarity-based hit expansion methods (25–27).

Overall, the three methods we created had better metrics than the work done by Schierz (19). The characteristics attributed by the Power MV tool increases the true positives rate but also increases the false positives rate. The classical methods have difficulties in obtaining high performance, being useful to use the Stratified Feature Selection. In the original paper(19), both bioassays, primary and confirmatory, we obtained high true positives rates. However, the study of the Bioassays combination was not as satisfactory, probably due to a big and unbalanced dataset, characteristic of Bioassays. The error associated with all methods can also be derived from the fact that the PubChem data is not cured. As the Milk-Way algorithm can select the features that contribute more to a ligand to be active and inactive, it is possible to segregate those ligands with more efficacy.

The free access of chemical and biological data in PubChem became a valuable resource for developing secondary databases and informatics tools that interact with PubChem. With this work, we believe that Milk-Way algorithm can be used as a virtual screening as a complement technique to the traditional approaches for virtual screening.

## Supporting information

Supplementary Material_Metrics

Supplementary Material_Values

Supplementary Material_Alphas

## LIST OF ABBREVIATIONS

AUC: Area under curve of Receiver Operating Characteristics Curve
HTS: High-Throughput Screening
LR: Logistic Regression
MW: Milk-Way algorithm
NB: Naïve Bayes
RF: Random Forest
SVM: Support Vector Machine

## ACKNOWLEDGEMENTS

To UCI Machine Learning Repository, by the available data and the possibility to use it (20).

## REFERENCES

1. Bellis LJ, Akhtar R, Al-Lazikani B, Atkinson F, Bento AP, Chambers J, et al. Collation and data-mining of literature bioactivity data for drug discovery. Biochemical Society transactions. 2011;39(5):1365–70.

2. Dahlin JL, Walters MA. The essential roles of chemistry in high-throughput screening triage. Future medicinal chemistry. 2014;6(11):1265–90.

3. Babaoglu K, Simeonov A, Irwin JJ, Nelson ME, Feng B, Thomas CJ, et al. Comprehensive mechanistic analysis of hits from high-throughput and docking screens against β-lactamase. Journal of medicinal chemistry. 2008;51(8):2502–11.

4. Ya-Di LI, Frenz CM, Mian-Hua C, Yu-Rong W, Feng-Juan LI, Cheng LUO, et al. Primary virtual and in vitro bioassay screening of natural inhibitors from flavonoids against COX-2. Chinese Journal of Natural Medicines. 2011;9(2):156–60.

5. Riniker S, Wang Y, Jenkins JL, Landrum GA. Using information from historical high-throughput screens to predict active compounds. Journal of chemical information and modeling. 2014;54(7):1880–91.

6. Sink R, Gobec S, Pecar S, Zega A. False positives in the early stages of drug discovery. Current medicinal chemistry. 2010;17(34):4231–55.

7. Wang Y, Suzek T, Zhang J, Wang J, He S, Cheng T, et al. PubChem bioassay: 2014 update. Nucleic acids research. 2013:gkt978.

8. Wang Y, Bolton E, Dracheva S, Karapetyan K, Shoemaker BA, Suzek TO, et al. An overview of the PubChem BioAssay resource. Nucleic acids research. 2010;38(suppl 1):D255–D66.

9. Kim S, Thiessen PA, Bolton EE, Chen J, Fu G, Gindulyte A, et al. PubChem substance and compound databases. Nucleic acids research. 2015:gkv951.

10. Bradley D. Dealing with a data dilemma. Nat Rev Drug Discov. 2008;7(8):632–3.

11. Ou-Yang S-s, Lu J-y, Kong X-q, Liang Z-j, Luo C, Jiang H. Computational drug discovery. Acta Pharmacologica Sinica. 2012;33(9):1131–40.

12. Koteluk O, Wartecki A, Mazurek S, Kołodziejczak I, Mackiewicz A. How Do Machines Learn? Artificial Intelligence as a New Era in Medicine. Journal of Personalized Medicine. 2021;11(1):32.

13. Han L, Wang Y, Bryant SH. Developing and validating predictive decision tree models from mining chemical structural fingerprints and high–throughput screening data in PubChem. BMC bioinformatics. 2008;9(1):1.

14. Riniker S, Fechner N, Landrum GA. Heterogeneous classifier fusion for ligand-based virtual screening: or, how decision making by committee can be a good thing. Journal of chemical information and modeling. 2013;53(11):2829–36.

15. Chen B, Wild D, Guha R. PubChem as a source of polypharmacology. Journal of chemical information and modeling. 2009;49(9):2044–55.

16. Paolini GV, Shapland RHB, van Hoorn WP, Mason JS, Hopkins AL. Global mapping of pharmacological space. Nature biotechnology. 2006;24(7):805–15.

17. Hao M, Wang Y, Bryant SH. An efficient algorithm coupled with synthetic minority over-sampling technique to classify imbalanced PubChem BioAssay data. Analytica chimica acta. 2014;806:117–27.

18. Hughes-Oliver JM, Brooks AD, Welch WJ, Khaledi MG, Hawkins D, Young SS, et al. ChemModLab: A web-based cheminformatics modeling laboratory. In silico biology. 2011;11(1, 2):61–81.

19. Schierz AC. Virtual screening of bioassay data. Journal of cheminformatics. 2009;1:21.

20. UCI Machine Learning Repository [Internet]. University of California, Irvine, School of Information and Computer Sciences. 2013. Available from: http://archive.ics.uci.edu/ml.

21. Figueiredo Vieira Leite C, Dos Santos MA, Silva Dos Santos LH, Fernando Leijôto L, Batista Mariano DC, Oliveira Rocha RE, inventors; Universidade Federal de Minas Gerais, assignee. Método de Triagem de compostos baseados em Regressão Logística Modificada. Brazil2019.

22. Leite CFV, Santos LHS, Leijôto LF, Mariano DCB, Rocha REO, Santos MAd. Milk-Way algorithm for ligand-based virtual screening: CDK2 case study. Trends in Developmental Biology. 2020;13.

23. Kurgan LA, Homaeian L. Prediction of structural classes for protein sequences and domains—Impact of prediction algorithms, sequence representation and homology, and test procedures on accuracy. Pattern Recognition. 2006;39(12):2323–43.

24. Dehzangi A, Paliwal K, Lyons J, Sharma A, Sattar A. Proposing a highly accurate protein structural class predictor using segmentation-based features. BMC Genomics. 2014;15 Suppl 1:S2.

25. Schürer SC, Vempati U, Smith R, Southern M, Lemmon V. BioAssay ontology annotations facilitate cross-analysis of diverse high-throughput screening data sets. Journal of biomolecular screening. 2011;16(4):415–26.

26. Clark AM, Litterman NK, Kranz JE, Gund P, Gregory K, Bunin BA. BioAssay templates for the semantic web. PeerJ Computer Science. 2016;2:e61.

27. Helal KY, Maciejewski M, Gregori-Puigjané E, Glick M, Wassermann AM. Public Domain HTS Fingerprints: Design and Evaluation of Compound Bioactivity Profiles from PubChem’s Bioassay Repository. Journal of Chemical Information and Modeling. 2016.

